# Benefits of combining individual and small group assessments as an instructional strategy

**DOI:** 10.1101/2024.10.21.619510

**Authors:** Gabriel Santos Arini, Israel de Souza Almeida, Bayardo Baptista Torres

## Abstract

Assessment is an essential curricular component despite being often seen only as a performance metric resource. However, its full potential can be harnessed if it is used as a formative instrument. The present study evaluated the benefits of combining individual and group assessments. Students’ perceptions of this type of strategy, assessed through a Likert scale questionnaire and semi-structured interviews, showed that students acknowledge the benefits of this procedure. We then carried out a dual assessment in an active learning environment. Students were given individual written tests and completed the same test immediately after, but now in a group. The average group scores were higher than the average individual scores, even for students who scored the highest within a group. Put together, these results indicate combining individual and group assessments can be an effective teaching tool in addition to simply measuring student performance.

## 1 Introduction

Assessment is a curricular component that branches out and touches virtually every other curriculum component. Furthermore, assessment is an essential part of learning and teaching. However, over time, assessment has lost its potential instructional richness. It is often underused, and the spectrum of its influence on the educational process has been reduced to its role as a tool for measuring the amount of information retained by the students. As a result, its diagnostic and “therapeutic” intrinsic attributes are undervalued. In addition, most assessment procedures share a weakness: the lack of associated measures to help low-achieving students.

To go beyond the limitations of conventional written tests, we have adopted and described a strategy that combines individual and group assessments, emphasizing peer discussion, metacognition [1], and teamwork [2]. The focus of this technique is to help students face (identify) and overcome their learning obstacles as soon as they appear. To reach this goal, individual and group assessments were designed and combined to act as instructional and evaluative tools that allow students to correct or reinforce their prior conceptions. This method was tested in a biochemistry course whose didactic strategy provided a suitable scenario for the investigation since students were familiar with small-group active learning tasks.

## 2 Procedures

### 2.1 Course description

This study was conducted in a 120-hour (for Chemistry students) or 180-hour (for Pharmaceutical Sciences students) introductory biochemistry course at the Department of Biochemistry of the University of São Paulo, Brazil. In each case, approximately 90 students were distributed into three medium-sized groups, each under the supervision of a faculty member. Each cohort was split into six small groups of up to 5 students. Classes took place two (Chemistry) or three (Pharmaceutical Sciences) times per week and lasted 4 hours each.

Following a quasi-classical sequence, the course content was organized into seven modules: Proteins, Enzymes, Glycolysis and Gluconeogenesis, Citric Acid Cycle and Fatty Acid Oxidation, Electron Transport Chain and Oxidative Phosphorylation, Glycogen Metabolism, and Amino Acid Metabolism.

The course replaces traditional lectures with small-group-directed study (DS) followed by group discussion sessions (GD). In the DS, small groups assemble to study, discuss, and answer low-order cognitive skill (LOCS) questions [3] presented in a study guide. To accomplish this task, the students can access books and software and receive guidance from faculty and teaching assistants (TA). After the completion of DS, GD takes place, when the small groups of each cohort are brought together, and students apply the information obtained in the DS to collaboratively solve high-order cognitive skills problems (HOCS) [3]. For each problem proposed for discussion, the group should reach a consensus, and any doubts should be clarified for everyone. Faculty members coordinate the discussions, watching for misconceptions and determining when the problem is resolved with the appropriate depth of understanding. DS and GD work synergistically to promote critical thinking, teamwork, and debating skills. The number of DS and GD sessions varied for each module.

### 2.2 Group assessment questionnaire

To address the effectiveness and students’ receptiveness to group tests, seven open-book tests, each at the end of a module, were administered to chemistry students to be solved by small groups of 4 or 5 students. The average of these tests made up 10% of the total grade. The remaining 90% was completed by the average of 4 open-book individual tests.

At the end of the term, students responded to a survey that addressed their perceptions and reactions to the assessment/learning strategy. A 5-point Likert scale (from strongly disagree to strongly agree) [4,5] was used to measure their level of agreement to 10 statements about the group assessment. Some redundancies in the items were intentionally designed to check if responses would not be question writing-related. Symmetrical counterparts of two items were introduced, as recommended for Cronbach’s reliability test. Semi-structured interviews were conducted to collect students’ opinions about the group assessment and its impact on their learning.

### 2.3 Dual assessment

Given the results obtained from the group assessment (see Results 3.1 and Figure 1), we designed a critical test to verify the learning efficacy of group assessment: a dual assessment. Following the DSs and GDs scheduled for each module, the students were required to individually complete a written open-book test consisting of 2-3 HOCS-level questions. Immediately after, the same test was completed in small groups, demanding students to discuss and collaboratively reach an answer for each question. The dual assessment method was introduced in the introductory biochemistry course for pharmaceutical sciences students.

**Figure 1.**
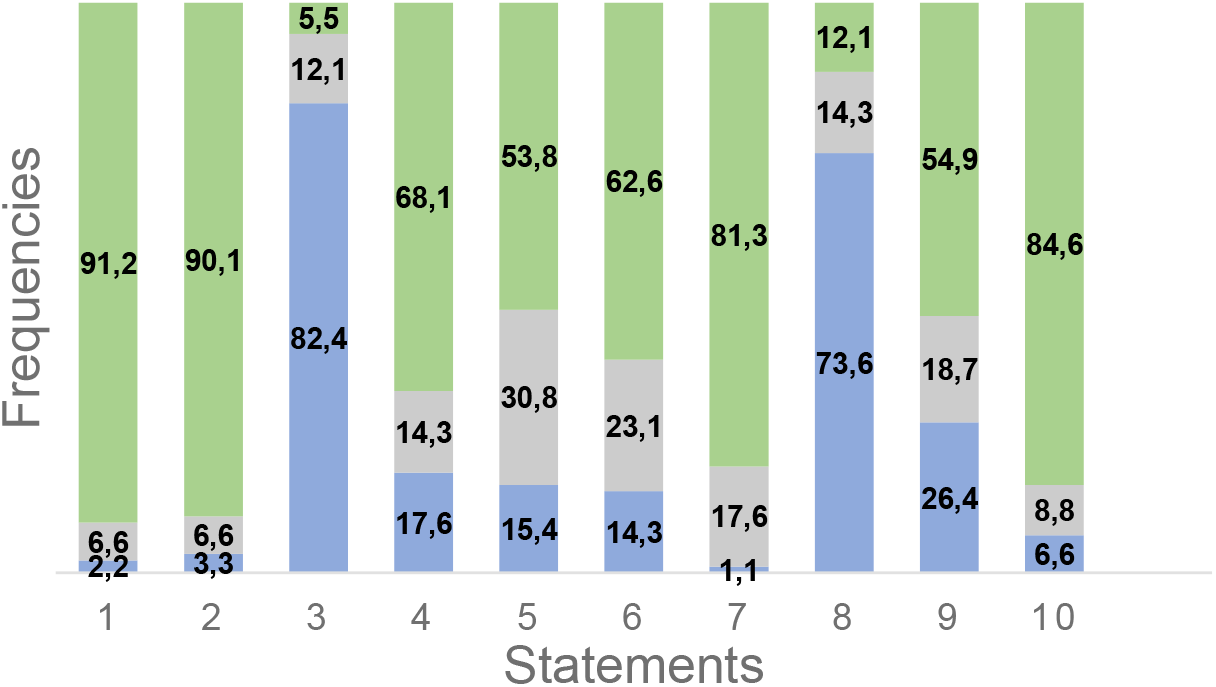
Student responses to the following statements: (1) The group assessment contributed to my learning. (2) The group discussion clarified the question when I initially did not know the answer. (3) The group assessments were formal and could be suppressed. (4) It would be nice if other courses adopted group quizzes. (5) Discussing group quizzes increased my interest in Biochemistry. (6) Group quizzes helped me develop my critical mind. (7) Group quizzes helped me develop my ability to analyze biochemical data. (8) Studying DS items replaces group quizzes. (9) Even if group quizzes were not graded, I would do them. (10) In general, all group members contributed to answering the questions. Green bars = Strongly agree + Agree; grey bars = neutral; blue bars = disagree + strongly disagree.

## 3 Results and Discussion

### 3.1 Receptiveness and efficiency of group assessment

Figure 1 shows the students’ level of agreement with the 10 statements in the questionnaire whose reliability was validated with a Cronbach’s alpha coefficient of 0.89 [6], well above the acceptable value of 0.7 commonly used to determine a questionnaire’s reliability [7]. The responses were 91, corresponding to 78.4% of the students enrolled in the course.

The high percentage of agreement with statements 1-2, 4-7, and 9 indicates that the students had a favorable opinion about the group assessment as a learning strategy. Coherently, 82.4 and 73.6% of the students disagreed with statements 3 and 8. Statements 6 and 7 are metacognitive and assess the development of skills extrapolating the limits of the course. Statement 9 deserves special attention: more than 50% of the students declared they would participate in the group assessments even if they were not graded, which stands for a recognition of this approach’s benefits.

To further explore the extent of the perceived benefits of the group test over traditional assessments, we conducted semi-structured interviews [8] with several key questions to address this issue. Representative queries and responses, translated from verbatim transcriptions, included the following:

1. If group assessments were not graded, would you continue to use them as a learning strategy? “I would, I would … I learned more in the group assessments than in the DSs and GDs”.
2. Did you notice any improvement in your performance due to group assessments? “Yes … it was during the group assessment that we discussed more in the group. There were many different questions. We had no way out and had to discuss until we got a consensual answer”.
3. If the group assessments were suppressed, would your data analysis skills still reach the same depth? “No, the group assessment deepened the concepts a lot and forced me to discuss them…”. Because of how we discussed, I do not think I would come to the same conclusions on my own.”
4. Did all group members actively participate in solving the problems in the group assessment? “Yes, everyone. All group members actively participated in solving the problem, and that is why it was worthwhile. “We were able to learn from each other. There were questions unknown to all of us, but through discussion, the group came up with an answer.

The latter individual declaration reinforces the answers to statement 10. Moreover, when asked to grade his/her participation in group assessments, the mean student’s score was 7.4 meaning that their opinion agreed with the group’s opinion.

In addition to the students’ perceptions of the strategy’s effectiveness, it is worth noting that faculty and TAs reported that the students were more engaged and active in taking the tests when they took the tests in groups.

### 3.2 Dual assessment

Frequent assessment and small-group active learning tasks play an important role in shaping a student’s approach to learning [9-12]. Group assessments are expected to improve learning as the debates required to reach a consensus can promote correction and broadening of previously acquired knowledge [13]. Our results in Figure 2 confirm this presupposition: mean group scores were significantly higher than individual mean scores regardless of faculty and maintained the same pattern over three consecutive years.

**Figure 2.**
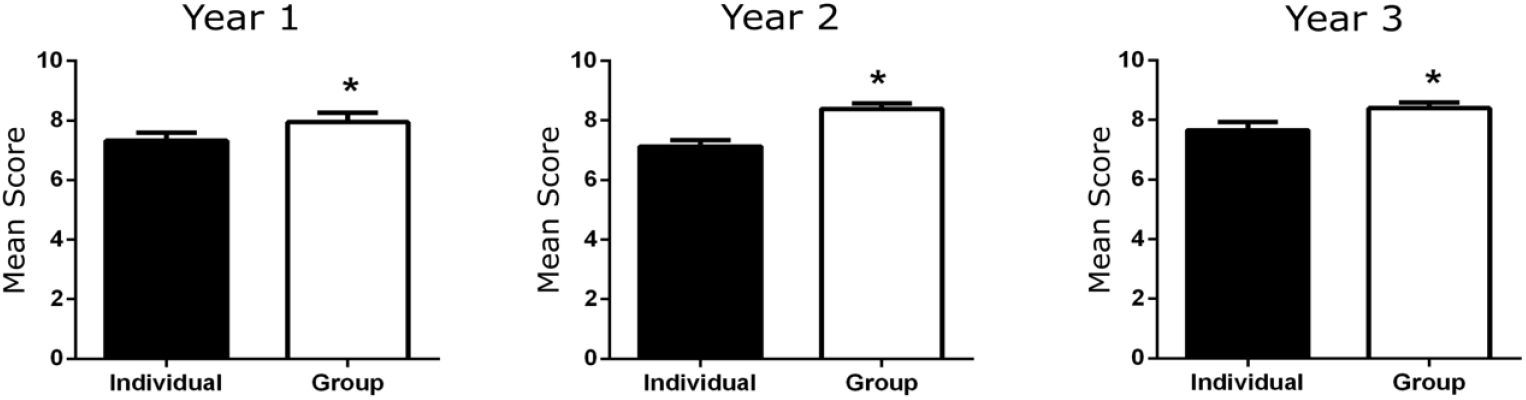
Individual’s and groups’ mean ± SEM scores. Number of students in each cohort: in year 1, Cohort A and B = 28; in year 2, Cohort A = 25; Cohort B = 28; and Cohort C = 30; in year 3, Cohort A = 21, Cohort B = 31, Cohort C = 24. Data was analyzed using paired Student’s t-test, *p<0.05 (individual vs. group). Black bars = Individuals’ mean scores; white bars = groups’ mean scores.

There are two alternative explanations for this outcome: it may result from gains in understanding during discussion or from the peer influence of knowledgeable students on their neighbors. The latter alternative can be discarded by comparing the best-performing students’ mean scores in each small group with the group’s mean score. This comparison is shown in Figure 3.

**Figure 3.**
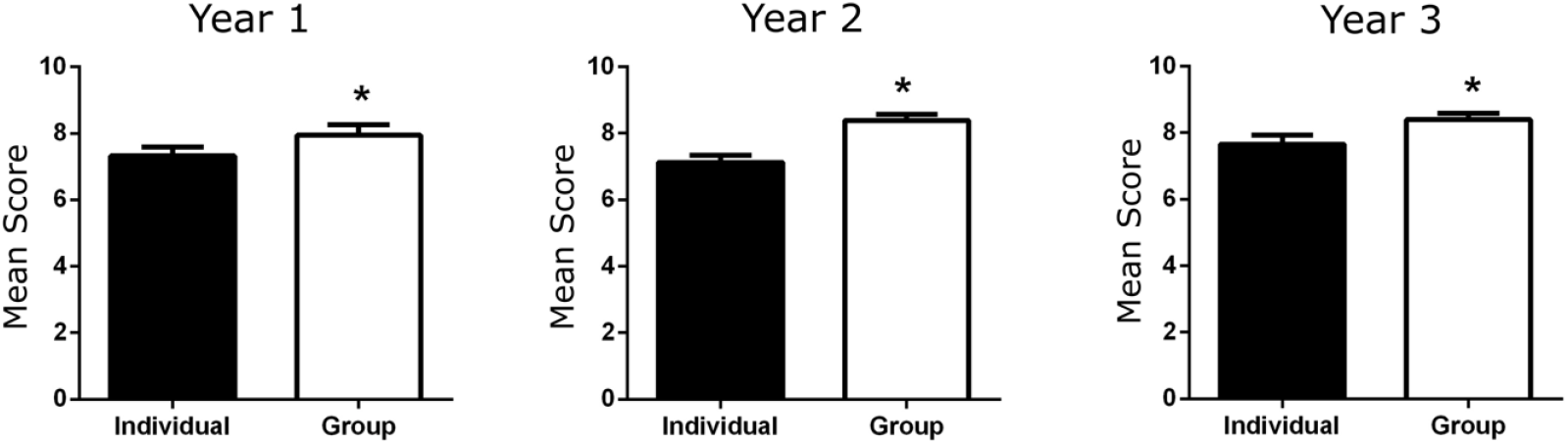
Best-performing students’ mean score in each cohort in each year and group mean score. Data was calculated using paired Student’s t-test mean ± SEM *p<0.05 (individual vs. group). The number of individuals was 11, 17, and 16 for year 1, year 2, and year 3, respectively. Black bars = individual mean score; white bars = groups’ mean scores.

In all cases studied, the group’s mean was higher than the score of the highest-scoring student. Therefore, his/her influence cannot account for the observed differences. Instead, this result must be interpreted as a learning gain generated by the group discussions since even the students with the highest scores benefited from the group discussion. The small difference between the individual and the group scores is expected since the best-performing students’ high scores leave little room for improvement, yet significant.

## 4 Final considerations

Our results showed that the average scores of students who participated in small-group assessments were consistently higher than those achieved individually, demonstrating the power and effectiveness of the dual assessment as an instructional tool. Among the perceived benefits is an improved ability to answer HOCS questions when comparing the scores of the highest-scoring students in each group to the group itself. One critical remark is that, in the student’s opinion, having someone in the group who knows the correct answer is unnecessary. The findings described here are consistent with the results of Smith et al. [13] from a different methodological set-up, namely a careful peer-instruction study in which group discussion precedes individual assessments. Group tests may, therefore, broaden the range of assessments by incorporating a valuable instructional attribute to traditional assessments.

